# An improved mouse model of sepsis based on intraperitoneal injections of the enriched culture of cecum slurry

**DOI:** 10.1101/2023.04.06.535817

**Authors:** Rajat Atre, Rahul Sharma, Alexander G Obukhov, Uzma Saqib, Sadiq Umar, Gajanan N Darwhekar, Mirza S Baig

## Abstract

Sepsis is a life-threatening clinical syndrome comprising multiorgan dysfunction caused by a disproportionate body immune response to infection that can lead to septic shock and death. The current sepsis model has certain limitations. The gold standard Cecal Ligation and Puncture (CLP) model is known for its high variability owing to the undefined extent of cecum ligation, unstandardized cecal content, and degree of puncture that may vary in different laboratories. Here, we present an improved, efficient intraperitoneal (i.p) injection-based cecal slurry culture method of sepsis. Using this novel approach, we determined the optimal polymicrobial concentration that is sufficient to cause sepsis in mice. We also proposed the enrichment of bacterial culture, allowing the development of either gram-negative or gram-positive bacteria-induced sepsis models. Since those enriched bacterial cultures can be stored in glycerol at −80°C, it gives the ethical advantage of avoiding animal sacrifice for each experiment and experimental reproducibility.

## Introduction

Sepsis is a life-threatening clinical condition resulting from the dysregulated immune response to an infection(Angus and van der Poll, 2013). Sepsis is characterized by unceasing inflammation that may lead to septic shock and death(Ayala and Chaudry, 1996; Delano and Ward, 2016; Gentile et al., 2012; Ward et al., 2008). Though advancements in techniques and strategies combating sepsis have been continuously worked on, sepsis has remained a consistent cause for high mortality(Steinhagen et al., 2021). According to an estimation of the World Health Organisation (WHO), over 30 million people worldwide are diagnosed with sepsis that results in ∼6 million deaths every year(Fleischmann et al., 2016). It has been reported that 50% of the patients who have survived a septic shock suffer from long lasting complications and are often repeatedly hospitalized(Huang et al., 2019). Furthermore, sepsis disproportionally affect the aged population of patients and especially those who present with weakened immune system(Juneja, 2012). Sepsis includes the failure of multiple organs which leads to devastating consequences. Despite the decades of intense research efforts, the pathophysiology of sepsis is not fully elucidated. However, the existing in vivo models of sepsis appear to be inadequate for studying this clinical condition(Walker, 2021).

To study the disease progression mechanism, the murine models of sepsis have been widely used by academic research groups and pharmaceutical companies involved in drug discovery and the development of potential therapeutic strategies against sepsis(Steinhagen et al., 2021). Though it has always been a topic of debate whether the animal models can fully mimic the sepsis condition of humans, these approaches still remain an integral part of the research eforts(Efron et al., 2015; Gentile et al., 2014b; Marshall et al., 2005; Seok et al., 2013). Despite various limitations, the conserved basic biology between the species in mammals and the ability to mimic the disease development makes the murine models an appropriate choice for such studies(Takao and Miyakawa, 2015).

Among the various methods used to induce sepsis, the CLP method is the gold standard approach. CLP involves induction of polymicrobial peritonitis by ligating the cecum surgically in anesthetized mice. Following this, puncture is made using a needle that results in the invasion of cecal contents into the abdominal cavity(Starr et al., 2014). Though it has been used for a long time, the approach has several limitations. The adversity of sepsis to be generated in the model is highly dependent on the severity of infection developed. Infection is influenced by various factors such as the bacterial population residing in the cecum, volume, and rate with which cecal contents get deployed into the abdominal cavity, compromising the reproducibility of this approach. Hence, despite being widely used, the method is incompetent when it comes to experiments involving animals with varying aforementioned factors.

Another method used for sepsis induction is the CASP approach, in which a plastic stent is inserted in anesthetized mice that allows the cecal contents from the colon to flow into the abdominal cavity(Maier et al., 2004; Zantl et al., 1998). The limitation of this method is the lack of information on the amount of microbial population being transferred from the colon into the abdominal cavity, bringing an uncertainty factor during the induction of sepsis.

Another alternative for sepsis induction is the CS method. In the CS sepsis model, mice are sacrificed, the cecal content is collected, and then stored as a frozen suspension at −80 °C. Intraperitoneal injection of the freshly thawed liquid suspension is intraperitonially given to other animals in order to induce polymicrobial sepsis(Gentile et al., 2014b, 2014a; Lang et al., 1983; Shrum et al., 2014; Wynn et al., 2007). The main drawback of the CS method is the need to sacrifice mice every time to gather the cecum slurry which contains variable microbial population, leading to prominent variability in the microbial populations among different preparations. Therefore, new approaches like slurry storage have been suggested in the recent past(Starr et al., 2014). But the lack of information regarding the microbial population being used remains a significant constraint of the recently proposed models for sepsis.

In the present study, we report the development of an improved method allowing reproducible and standardized sepsis induction. Our novel approach is based on the intraperitoneal injection of a known amount of enriched polymicrobial population derived from cecum slurry and propagated in LB media until a desired bacterial density, precisely measured by optical density (OD). Herein we propose the storage of the cultured poly microbiota in glycerol stocks at −80°C, allowing us to maintain a constant bacterial population among multiple experiments (Fig.1). The proposed model gives an ethical advantage over previous methods because it avoids unnecessary animal sacrifice for preparing the cecum slurry for each performed experiment. Remarkably, we propose an enrichment approach (Fig. 3) that permits us to selectively grow Gram-positive or Gram-negative bacterial cultures from cecum slurry for inducing standardized sepsis models.

## Results

### 1. Effect of different bacterial concentrations on survival of mice

To determine the effect of bacterial concentrations on the mortality of mice presenting with experimental sepsis, we divided 44 mice into 5 groups (A, B, C, D, and E), with each group containing 8-10 mice (2 extra mice in group C and D for tissue collection). The mice were intraperitonially injected with either 2×10^9^ CFU/ml (group A), 1×10^9^ CFU/ml (group B), 0.5×10^9^ CFU/ml (group C), or 0.1×10^9^ CFU/ml (group D) bacterial suspension, respectively, whereas the group E was kept as control (Fig. 1). To achieve the aforementioned bacterial concentrations, we used a bacterial culture with an optical density of 1OD (determined at 600 nm) and transferred from it 2 ml, 1 ml, 0.5 ml, and 0.1 ml into four sterile individual centrifuge tubes. It has been reported that an 1OD bacterial culture has a bacterial concentration of about 10^9^ CFU in 1 ml of culture media(Stevenson et al., 2016). After centrifugation, each pellet was dissolved in 500 μl 1X PBS and intraperitonially injected per mouse. Mice were monitored after every 2 hrs. We found 80% mortality at 6 hours in group A, 50% mortality at 6 hours in group B, 50% mortality at 16 hours in group C, and only 20% mortality at 20 hours in group D. 100% mortality was observed in group A with the maximum bacterial concentration after 16 hours post injection. Conversely, in the group D with the minimum concentration only 20% mortality was observed at 20 hours post injection, and all survivor mice showed fast recovery (Fig. 1). When closely inspected, all mice in group A, B, and C developed white eyes and loss of vision after 12 hrs. indicating severe sepsis condition(Grimm and Willmann, 2012). The mouse eyes were unchanged in group D, which indicates the insufficient polymicrobial concentration to cause sever sepsis. The whole experiment was replicated using the same glycerol stock of the polymicrobial cultures to validate the results.

**Figure 1.**
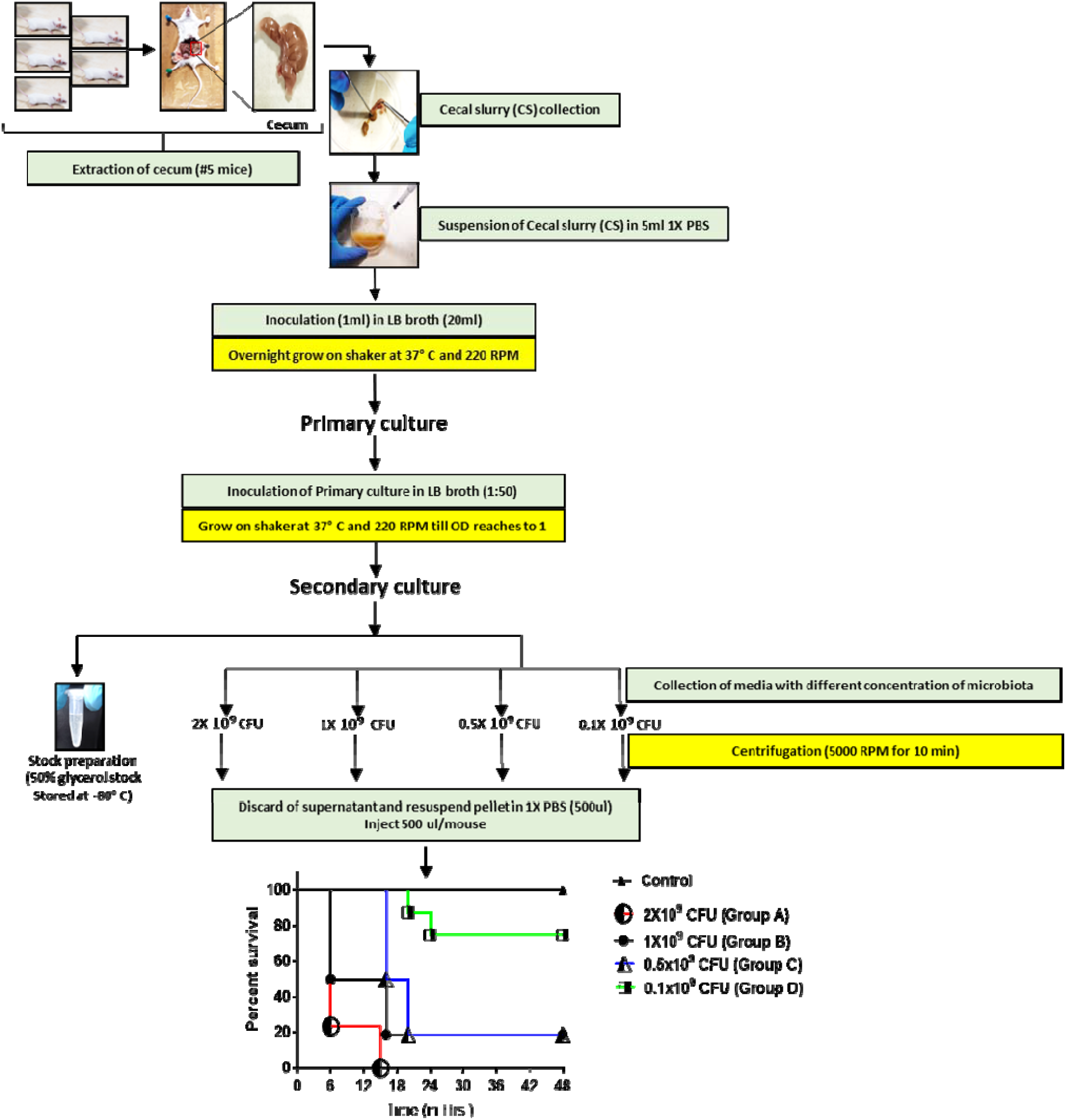
Protocol for the improved caecal slurry method of sepsis and the survival curve in mice treated with 2×10^9^ CFU/ml, 1×10^9^ CFU/ml, 0.5×10^9^ CFU/ml, and 01×10^9^ CFU/ml (8 mice per group).

### 2. Effect of different bacterial concentration on different organs of mice

To see the effect of different bacterial concentration, we collected tissue samples of the lungs, liver and kidney in group C and D to compare the organ damage and to quantify inflammation between the maximum tolerated concentration (group C) and the insufficient polymicrobial concentration (group D) to cause sepsis. All the samples were collected after 10 hours after injection. We dissected two mice from each group and collected the indicated organs in 4% formaldehyde after a quick wash with 1X PBS. The samples were then sectioned and subjected to H&E staining. We compared the H&E-stained lung tissue of Group D mice with the untreated one (Fig. 2A) and observed more infiltration of immune cells (depicted by black arrow) and increment in necrosis (depicted by red arrow) in the septum, indicating inflammation and tissue damage. Consistently, the group C lung tissue had more infiltrated immune cells than the group D. We also quantified the number of cells infiltrating in the alveolar septum (Fig. 2B) and the vacant space between the septum (Fig.2E) by using ImageJ bundled with 64-bit Java 8 and 536×389 pixels images. We found that the degree of immune cell infiltration correlated well with the polymicrobial concentration, and the vacant space was decreased in accordance with it, clearly indicating that sepsis was more significant with increasing inflammation. The alveoli were damaged more in group C compared to group D, further confirming that the severity of sepsis increased in group C. We next compared the liver tissue (Fig. 2A). We found more immune cells as well as larger necrosis affected areas in group C than in group D, again indicating that inflammation and tissue damage was increased with high polymicrobial concentration. We observed a similar pattern in the kidneys (Fig. 2A). We also counted the number of infiltrating immune cells and the percent of necrotic area in the liver (Fig. 2C and 2F) and kidney (Fig. 2D and 2G) and found that the number and necrotic area increased. In short, all the three organs had increased inflammation and exhibited the signs of multi-organ damage that likely led to sepsis and then further to septic shock, resulting in animal death.

**Figure 2.**
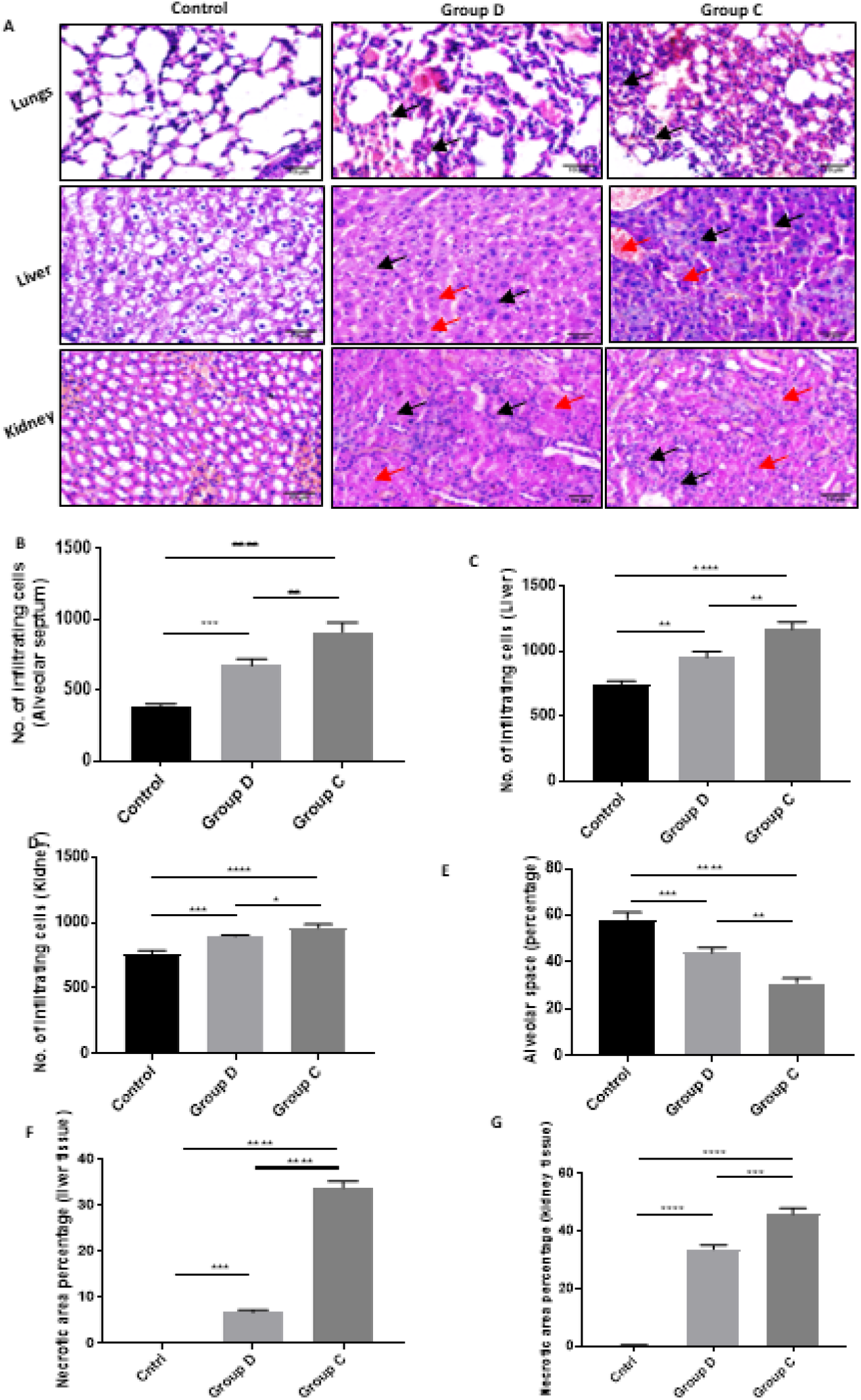
Effect of caecal slurry polymicrobial concentration on mice. (A) H&E-stained sections of the lung, liver and kidney tissue at 40X (10 μm scale bars). The samples were collected in mice without treatment, as well as in mice treated with 0.5×10^9^ CFU/ml and 0.1×10^9^ CFU/ml of polymicrobial cultures (black arrow and red arrow depicts infiltration of immune cells and necrotic area, respectively). (B)-(D) shown are the numbers of immune cells infiltrating in the alveolar septum, liver, and kidney. (E) shown are the alveolar space percentage between the alveolar septum. (F)-(G) are showing the necrotic area in terms of percentage in liver and kidney, respectively. The whole experiment was replicated with the 50% glycerol stock of polymicrobial cultures. All data are representative of two independent experiments; all data are presented as mean ± SD. P Significance was determined using One-way ANOVA test. **P ≤ 0.01; *P ≤ 0.01.

**Figure 3:**
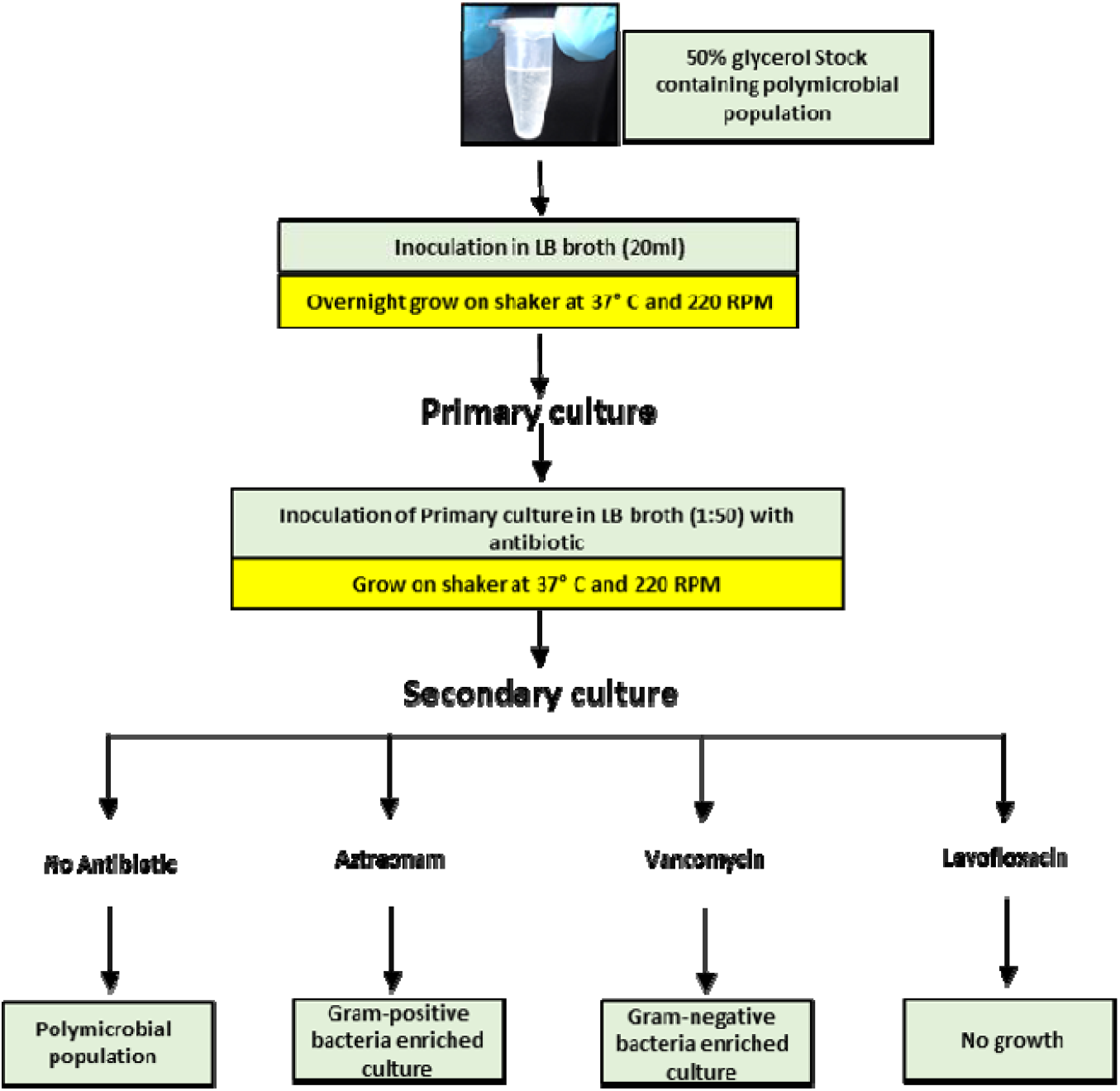
Enrichment protocol - Glycerol stock was cultured with respective antibiotics to generate specific bacterial populations.

### 3. Inducing gram-negative and gram-positive sepsis by enriching bacterial cultures

The above experiments were performed to induce polymicrobial sepsis. We next used the same approach but introduced the enriched bacterial cultures to induce gram-negative or gram-positive bacterial sepsis. We employed two antibiotics, aztreonam and vancomycin, which can selectively target gram-negative or gram-positive bacteria, respectively. Additionally, we used a broad-spectrum antibiotic levofloxacin to determine if there was any growth other than bacteria. We added 5 ml LB media in each of five 15 ml tubes and supplemented the media with respective antibiotics (20 μg/ml) and labelled the tubes as follows: tube A (Blank), tube B (LB + Inoculum), tube C (LB + Aztreonam + Inoculum), tube D (LB + Vancomycin + Inoculum), and tube E (LB + Levofloxacin + Inoculum). The media containing specific antibiotic was inoculated with 1:50 part of primary culture (1.5 OD) and incubated at 37 °C for 3 hours on a shaker with a rate of 220 rotation per minute. The tubes were then placed on ice and OD was measured using an UV spectrophotometer at 600 nm to determine the growth pattern of bacteria under different conditions. The OD of culture media containing no antibiotics was 0.75 while Aztreonam, vancomycin and levofloxacin enriched cultures had a value of 0.41, 0.7, and 0.05 OD, respectively (Fig 4A), indicating that the broad-spectrum antibiotic levofloxacin can kill all type of bacteria and that there were no fungi in the cecal slurry. Notably, the gram-positive bacteria grew at lower rate than the gram-negative bacteria. For further validation, we also performed spreading of polymicrobial culture on agar plates with no antibiotics (Fig. 4C-a), aztreonam (Fig. 4C-b), vancomycin (Fig. 4C-c), or levofloxacin (Fig. 4C-d). The results showed the same pattern of growth as observed by measuring the OD values at 600 nm. To confirm the action of respective antibiotics, we next performed gram staining by taking 20 μl of suspension from each tube and making smears on slides. We utilized the HIMEDIA-K001-1KT kit for gram staining and analysed the stained smears using the (Brand) microscope at 40X. In cecal slurry, there was growth of all type of bacteria that can be visualized by violet and pink stains (Fig 4B-a). There were more gram-positive bacilli bacteria in tube C identified by the crystal violet (Fig. 4B-b) and further the strings of gram-positive bacilli, which cannot be formed by gram negative bacilli (https://tmedweb.tulane.edu/pharmwiki/doku.php/bacteria_101_cell_walls_gram_staining_common_pathogens), can also be observed. Accordingly, there were more gram-negative cocci and bacilli in Tube D (Fig 4B-c). As expected, there was no growth of any microbial population in Tube E (Fig. 4B-d). For future experiments, the enriched culture can be stored at −80°C as 50% glycerol stocks or can be used for inducing gram-negative or gram-positive sepsis by using the respective cultures at 1 OD and injecting the mice with the desired bacterial concentrations. Furthermore, the cultures may be propagated to amplify specific bacterial population.

**Figure 4.**
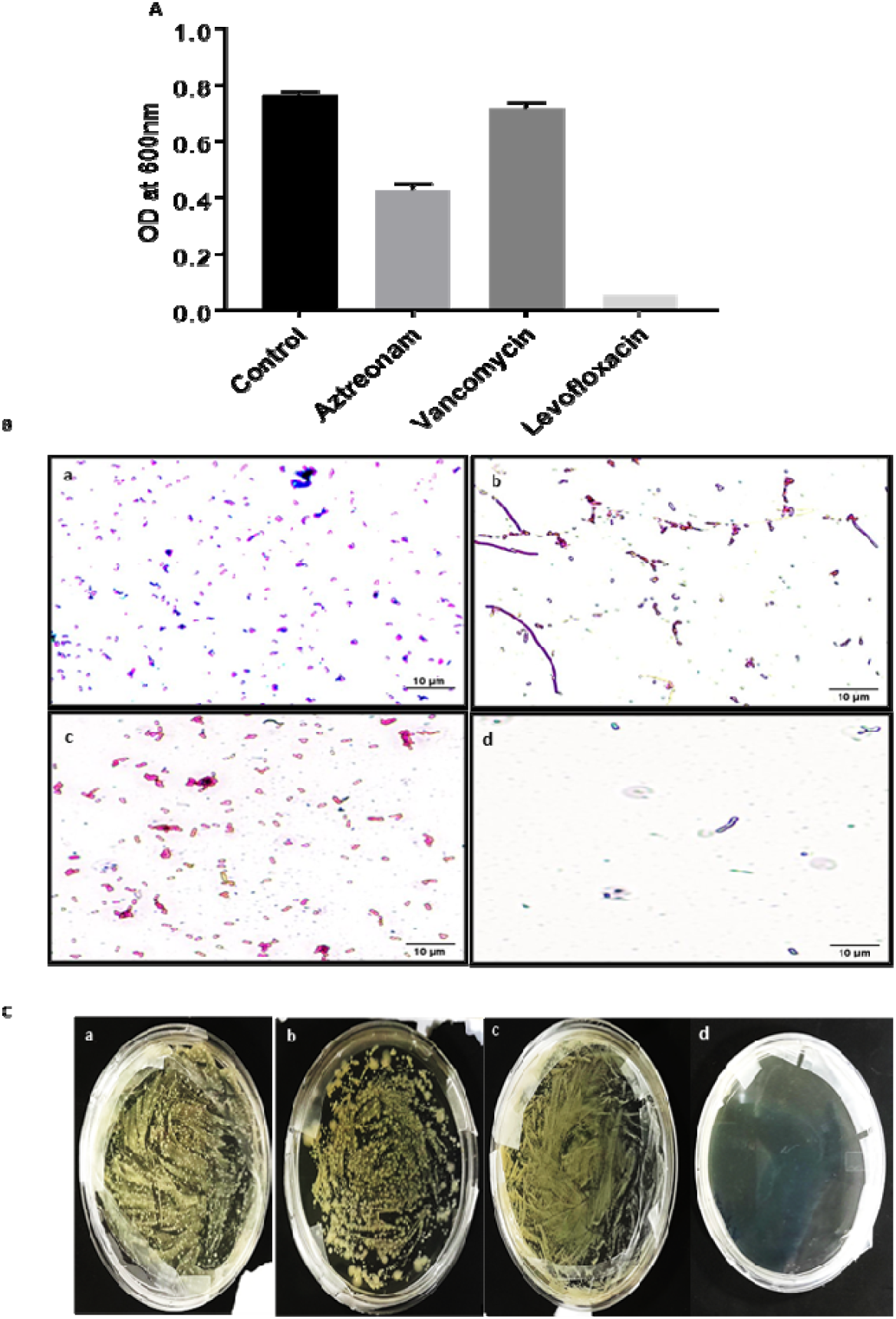
Culturing cecal slurry polymicrobiota in media enriched with antibiotics (20 ug/ml) targeting gram –ve and +ve bacteria. (A) The bar graph depicts the growth rate of bacteria under different conditions, where Aztreonam, vancomycin, and levofloxacin targets gram –ve bacteria, gram +ve and both, respectively. Levofloxacin was used to determine whether there are any other microbes beside bacteria in the cecal slurry. The OD was taken after 3 hours and the error bar shows the OD value after 3.30 hours. (B) The gram-stained images at 40X, where (a) the culture without antibiotics, (b) aztreonam treated culture, (c) vancomycin treated culture, and (d) levofloxacin treated culture. (C) The growth patterns with different antibiotics. LB culture media was spread on LA with (a) no antibiotics (b) aztreonam (c) vancomycin (d) levofloxacin.

## Discussion

Sepsis is a common clinical condition that is associated with a high morbidity and mortality throughout the globe(Juneja, 2012). This life-threatening syndrome may occur due to the unceasing exaggerated inflammatory response along with an immunosuppressed state(Ayala and Chaudry, 1996; Delano and Ward, 2016; Gentile et al., 2012; Ward et al., 2008). Though many advancements have been made in the direction of finding concrete solutions to treat sepsis, it still remains one of the most serious clinical problems. It is very important to understand the pathophysiology of the disease to design effective therapeutic strategies against this condition. Considering the complexity and the involvement of various organs in the biology of sepsis, it is the immediate requirement to develop better therapies against this multi organ dysfunction condition.

Animal sepsis models are widely used to study the pathophysiology of sepsis. However, it is crucial to identify the right model which can be reproducible and capable of mirroring both cellular and molecular aspects of sepsis, such as observed in humans. Several approaches have been developed in the past to study sepsis. These include the CLP(Dejager et al., 2011), CASP(Walker, 2021), CS(Gentile et al., 2014b, 2014a; Lang et al., 1983; Shrum et al., 2014; Wynn et al., 2007) methods and few others.

The CLP approach is considered as the “Gold standard” to develop and replicate the human condition of polymicrobial infection in animals(Dejager et al., 2011). Despite its wide acceptance, it has several shortcomings. This model is complicated due to high variability in terms of size and shape of cecum that ultimately affects sepsis severity. Similarly, such models as CASP(Maier et al., 2004; Zantl et al., 1998) and CS(Gentile et al., 2014b, 2014a; Lang et al., 1983; Shrum et al., 2014; Wynn et al., 2007) that are also aimed to mirror sepsis conditions recapitulating the human state have limitations. These models require the sacrifice of additional animals for every single new experiment. Furthermore, these approaches do not provide a sufficient insight on the microbial flora composition that is delivered into the abdominal cavity from the cecum, bringing the uncertainty factor in the induction of sepsis model. In the CS model, the slurry is prepared fresh every time(Starr et al., 2014), thus adding a high amount of variability every time the experiment is performed. Though a new method was recently developed where storage option has been proposed, making an alternative to animal sacrifice for each experiment, the approach still lacks the reproducibility of sepsis conditions among different experiments and again lacks information on the composition of microbial population. Hence, a better and improved approach is required to tackle the obstacles being faced in the development of sepsis models.

Herein we propose a novel and efficient method for inducing polymicrobial sepsis in animals. Our protocol is based on the intraperitoneal injections of defined bacterial cultures with known polymicrobial population. In our experiments, we compared five animal groups, including a control group. Each treatment group received varying doses of polymicrobial bacterial load. Before animal injection, the polymicrobial cultures were grown in the standard LB medium that is a standardizable approach, allowing sepsis induction with varied titrated inoculum dosages in order to develop the diseased state with varying severity as per the requirement. The results from the survival curves depict that the groups injected with higher CFU had early mortality and the mortality rate decreased with the decreases in CFU. This can prove to be helpful in order to regulate and plan the experiments to better understand the biology and various factors involved at varying stages of sepsis.

Other methods required the cecum slurry to be freshly prepared for every new experiment; new advancements have made it possible to store frozen cecum slurry suspensions(Starr et al., 2014). In our study, we make a step forward and propose the storage of the artificially propagated polymicrobial culture derived from cecal slurry that can be stored at -80°C in 50% glycerol for a very long period. In our experiments, similar results were obtained in replicate experiments using the frozen bacterial stocks indicating that the frozen bacteria preserved viability.

As sepsis is a multiorgan dysfunction condition, we extracted different organs, which are supposedly affected by sepsis, to see the effect of the varied microbial population used for sepsis induction. The lungs, liver, and kidneys were analysed for infiltration of immune cells to compare the extent of inflammation using H&E-staining. The severity of sepsis correlated with the survival results, confirming our ability to successfully monitor disease progression. Importantly, our approach provides ethically benefits as it does not require to sacrifice animals for every new experiment.

In this study, we also focused on overcoming the inconsistency of previously proposed methods to standardize the bacterial inoculum(Starr et al., 2014). We put forward a culture enrichment strategy that permits us to grow a specific population i.e.; gram-negative or gram-positive bacteria. Uniquely, we used special antibiotics to harvest these bacterial species’ specific cultures and validated the results via gram staining. This enrichment method could give the researchers an effective and efficient way to generate specific cultures in order to investigate various disease models which require a defined microbial population. Moreover, without the usage of any special antibiotics one can just grow a polymicrobial bacterial population from the same initial bacterial culture stored at -80° C as glycerol stocks. Thus, avoiding the variability associated with previous approaches which utilized undefined cecum slurry preparations for sepsis induction.

In sum, we propose an OD based polymicrobial sepsis model that tends to overcome several limitations of previous sepsis models. The defined polymicrobial cultures can be stored at −80° C as glycerol stocks for prolonged durations and later propagated as needed. The major advantages of our protocol are 1) Ability to standardize the bacterial inoculum, 2) Ethically suitable as it limits unnecessary sacrifice of animals, 3) Effective and technically sound method, and 4) Enrichment approach allowing to culture specific defined polymicrobial populations. These advances would definitely help further our understanding of the pathophysiology of sepsis and may contribute to saving lives of affected patients.

## Methods

### Animal studies

4–5-month-old male and female mice in the ratio of 3:2, each weighing 30 g, were obtained from the College of Veterinary Science and Animal husbandry, Mhow, Indore, Madhya Pradesh, India. All mice were kept at 25°-28°C and a 14:10 h of light and dark cycle with 24×7 available drinking water and chow. Mice were acclimated for 7 days prior to experiment. This animal study was approved by the Institutional Animal Ethics Committee (IAEC) of Acropolis Institute of Pharmaceutical Education and Research and conducted in accordance with the policies of Committee for the Purpose of Control and Supervision of Experiments on Animals (CPCSEA), Govt. of India.

### Preparation of cecum slurry

Five (3 male + 2 female) 4–5-month-old mice were used for initial cecal slurry collection. They were first sacrificed by cervical dislocation under anaesthesia (Chloroform) and the whole cecum was dissected with sterile forceps and scissors. By using forceps and spatula, the entire cecal content was collected in a sterile petri dish and the whole content was then mixed with 5 ml sterile 1X PBS by continuous pipetting.

### Culture and stock preparation

1ml of cecal slurry resuspended in 1X PBS was inoculated in 50 ml Luria Broth (HIMEDIA M575-500G) and was cultured overnight at 37°C on a shaker (220 RPM). The culture was then filtered (70 μm mesh strainer) to remove slurry particles. 1/60th part of that 1° culture (OD=1.5) was then inoculated in 50 ml LB media (2° culture). By utilizing this inoculum volume, the OD reached to 1 after 3 hours of incubation. OD was measured using UV spectrophotometer (PerkinElmer UV/VIS Spectrometer Lambda 35) at 600 nm. Different volumes of the culture media were then used for preparing injections with different bacterial concentration and the remaining media was used for preparing cryovials of 50% glycerol stock which were then stored at −80°C. For enrichment of cultures with a particular type of bacteria, the following specific antibiotics (20 ug/ml) were used: Aztreonam to kill Gram negative bacteria, and Vancomycin to kill Gram positive bacteria. Antibiotics were added to the secondary culture. To check the growth rate of bacteria in these antibiotics rich media, OD was measured at 600 nm. To validate the growth rate, spreading with primary culture was additionally done on agar plates containing Luria Agar (HIMEDIA M557-500G) and respective antibiotics. These were then observed after overnight incubation at 37°C. Gram staining (see below) was performed to confirm the type of microbiota in polymicrobial population. This enrichment culture method was replicated with both fresh cecal slurry and the 2 months stored 50% glycerol stock.

### Preparation samples for injections

Injection samples were prepared for four groups of 8 mice each, with different bacterial concentrations that were 2×10^9^ CFU/ml, 1×10^9^ CFU/ml, 0.5×10^9^ CFU/ml, and 0.1×10^9^ CFU/ml per mice, by taking 2ml, 1ml, 0.5ml, and 0.1ml of culture media (1 OD) per mouse, respectively. The media was then centrifuged at 6000 G at 4°C and then the pellet containing bacteria was resuspended in sterile 1X PBS and injected at 500 μl per mouse.

### H&E staining

Haematoxylin and eosin stain was used for staining nuclei (blue) and cytoplasm/extracellular space (pink). The tissue samples preserved in 4% formaldehyde were used for mounting and H&E staining. The prepared slides were then observed using an Invitrogen EVOS M5000 Cell Imaging System, and the images were taken at 40X. The alveolar space and the infiltrating immune cells were counted by using ImageJ.

### Gram staining

To validate the enriched culture, whether it is gram positive or negative bacteria, a simple gram staining technique was performed by using gram staining kit (HIMEDIA-K001-1KT), where crystal violet was used to stain gram positive while safranin was used to stain gram negative bacteria. The culture media was used immediately after measuring the OD.

### Mortality analysis

Mice were injected with different polymicrobial concentration and were monitored for up to 48 hours post injection. The Kaplan-Meier curve was used for graphical presentation, where Y-axis represented percent survival. The tissue samples of the lungs, liver and kidneys were collected 10 hours post injection in animals from two groups (2 mice/group). These were preserved in 4% formaldehyde and were analysed after H&E staining. Each experiment was performed twice, and the survival curve was constructed by the data acquired in each group.

### Statistical analysis

Statistical analysis and visualization of data was performed using GraphPad PRISM 7.0.4.216 (© 1992-2017 GraphPad Software. lnc). All data are representative of two independent experiments; all data are presented as mean ± SD. P Significance was determined using One-way ANOVA test. **P ≤ 0.01; *P ≤ 0.01.

## Acknowledgment

The authors gratefully acknowledge the Indian Institute of Technology Indore (IITI) for providing facilities and other support. The Acropolis Institute of Pharmaceutical Education and Research (AIPER, Indore) for allowing us to use its animal housing facility. We also thank the Choithram Hospital and Research Centre (CHRC) Indore for lung tissue sectioning and staining.

## Author Contributions

Conceptualization and investigation: MSB; Experiments: RS and RA; Writing (Original Draft): MSB, RS, and RA; Reviewing and editing AGO, US, SU, and GND. All authors have read and agreed to the published version of the manuscript.

## Data Availability Statement

The original contributions presented in the study are included in the article. Further inquiries can be directed to the corresponding authors.

## Ethics Statement

All animal studies described in this paper were performed in accordance with the animal protocol which was approved by the Institutional Animal Ethics Committee (IAEC) of Acropolis Institute of Pharmaceutical Education and Research and following the policies of the Committee for Control and Supervision of Experiments on Animals (CPCSEA), Government of India.

## References

Angus DC, van der Poll T. 2013. Severe Sepsis and Septic Shock. N Engl J Med 369:840– 851. doi:10.1056/NEJMra1208623

Ayala A, Chaudry IH. 1996. Immune dysfunction in murine polymicrobial sepsis: mediators, macrophages, lymphocytes and apoptosis. Shock 6 Suppl 1:S27–38.

Dejager L, Pinheiro I, Dejonckheere E, Libert C. 2011. Cecal ligation and puncture: the gold standard model for polymicrobial sepsis? Trends in Microbiology 19:198–208. doi:10.1016/j.tim.2011.01.001

Delano MJ, Ward PA. 2016. The immune system’s role in sepsis progression, resolution, and long-term outcome. Immunol Rev 274:330–353. doi:10.1111/imr.12499

Efron PA, Mohr AM, Moore FA, Moldawer LL. 2015. The future of murine sepsis and trauma research models. Journal of Leukocyte Biology 98:945–952. doi:10.1189/jlb.5MR0315-127R

Fleischmann C, Scherag A, Adhikari NKJ, Hartog CS, Tsaganos T, Schlattmann P, Angus DC, Reinhart K. 2016. Assessment of Global Incidence and Mortality of Hospital-treated Sepsis. Current Estimates and Limitations. Am J Respir Crit Care Med 193:259–272. doi:10.1164/rccm.201504-0781OC

Gentile LF, Cuenca AG, Efron PA, Ang D, Bihorac A, McKinley BA, Moldawer LL, Moore FA. 2012. Persistent inflammation and immunosuppression: A common syndrome and new horizon for surgical intensive care. Journal of Trauma and Acute Care Surgery 72:1491–1501. doi:10.1097/TA.0b013e318256e000

Gentile LF, Nacionales DC, Lopez MC, Vanzant E, Cuenca A, Cuenca AG, Ungaro R, Szpila BE, Larson S, Joseph A, Moore FA, Leeuwenburgh C, Baker HV, Moldawer LL, Efron PA. 2014a. Protective Immunity and Defects in the Neonatal and Elderly Immune Response to Sepsis. The Journal of Immunology 192:3156–3165. doi:10.4049/jimmunol.1301726

Gentile LF, Nacionales DC, Lopez MC, Vanzant E, Cuenca A, Szpila BE, Cuenca AG, Joseph A, Moore FA, Leeuwenburgh C, Baker HV, Moldawer LL, Efron PA. 2014b. Host Responses to Sepsis Vary in Different Low-Lethality Murine Models. PLoS ONE 9:e94404. doi:10.1371/journal.pone.0094404

Grimm C, Willmann G. 2012. Hypoxia in the Eye: A Two-Sided Coin. High Altitude Medicine & Biology 13:169–175. doi:10.1089/ham.2012.1031

Huang CY, Daniels R, Lembo A, Hartog C, O’Brien J, Heymann T, Reinhart K, Nguyen HB, Sepsis Survivors Engagement Project (SSEP). 2019. Life after sepsis: an international survey of survivors to understand the post-sepsis syndrome. International Journal for Quality in Health Care 31:191–198. doi:10.1093/intqhc/mzy137

Juneja D. 2012. Severe sepsis and septic shock in the elderly: An overview. WJCCM 1:23. doi:10.5492/wjccm.v1.i1.23

Lang CH, Bagby GJ, Bornside GH, Vial LJ, Spitzer JJ. 1983. Sustained hypermetabolic sepsis in rats: Characterization of the model. Journal of Surgical Research 35:201– 210. doi:10.1016/S0022-4804(83)80005-5

Maier S, Traeger T, Entleutner M, Westerholt A, Kleist B, Hser N, Holzmann B, Stier A, Pfeffer K, Heidecke C-D. 2004. CECAL LIGATION AND PUNCTURE VERSUS COLON ASCENDENS STENT PERITONITIS: TWO DISTINCT ANIMAL MODELS FOR POLYMICROBIAL SEPSIS: Shock 21:505–512. doi:10.1097/01.shk.0000126906.52367.dd

Marshall JC, Deitch E, Moldawer LL, Opal S, Redl H, Poll T van der. 2005. PRECLINICAL MODELS OF SHOCK AND SEPSIS: WHAT CAN THEY TELL US? Shock 24:1–6. doi:10.1097/01.shk.0000191383.34066.4b

Seok J, Warren HS, Cuenca AG, Mindrinos MN, Baker HV, Xu W, Richards DR, McDonald-Smith GP, Gao H, Hennessy L, Finnerty CC, López CM, Honari S, Moore EE, Minei JP, Cuschieri J, Bankey PE, Johnson JL, Sperry J, Nathens AB, Billiar TR, West MA, Jeschke MG, Klein MB, Gamelli RL, Gibran NS, Brownstein BH, Miller-Graziano C, Calvano SE, Mason PH, Cobb JP, Rahme LG, Lowry SF, Maier RV, Moldawer LL, Herndon DN, Davis RW, Xiao W, Tompkins RG, the Inflammation and Host Response to Injury, Large Scale Collaborative Research Program, Abouhamze A, Balis UGJ, Camp DG, D. AK, Harbrecht BG, Hayden DL, Kaushal A, O’Keefe GE, Kotz KT, Qian W, Schoenfeld DA, Shapiro MB, Silver GM, Smith RD, Storey JD, Tibshirani R, Toner M, Wilhelmy J, Wispelwey B, Wong WH. 2013. Genomic responses in mouse models poorly mimic human inflammatory diseases. Proc Natl Acad Sci USA 110:3507–3512. doi:10.1073/pnas.1222878110

Shrum B, Anantha RV, Xu SX, Donnelly M, Haeryfar SM, McCormick JK, Mele T. 2014. A robust scoring system to evaluate sepsis severity in an animal model. BMC Res Notes 7:233. doi:10.1186/1756-0500-7-233

Starr ME, Steele AM, Saito M, Hacker BJ, Evers BM, Saito H. 2014. A New Cecal Slurry Preparation Protocol with Improved Long-Term Reproducibility for Animal Models of Sepsis. PLoS ONE 9:e115705. doi:10.1371/journal.pone.0115705

Steinhagen F, Hilbert T, Cramer N, Senzig S, Parcina M, Bode C, Boehm O, Frede S, Klaschik S. 2021. Development of a minimal invasive and controllable murine model to study polymicrobial abdominal sepsis. All Life 14:265–276. doi:10.1080/26895293.2021.1909663

Stevenson K, McVey AF, Clark IBN, Swain PS, Pilizota T. 2016. General calibration of microbial growth in microplate readers. Sci Rep 6:38828. doi:10.1038/srep38828

Takao K, Miyakawa T. 2015. Genomic responses in mouse models greatly mimic human inflammatory diseases. Proc Natl Acad Sci USA 112:1167–1172. doi:10.1073/pnas.1401965111

Walker WE, editor. 2021. Sepsis: Methods and Protocols, Methods in Molecular Biology. New York, NY: Springer US. doi:10.1007/978-1-0716-1488-4

Ward NS, Casserly B, Ayala A. 2008. The Compensatory Anti-inflammatory Response Syndrome (CARS) in Critically Ill Patients. Clinics in Chest Medicine 29:617–625. doi:10.1016/j.ccm.2008.06.010

Wynn JL, Scumpia PO, Delano MJ, O’Malley KA, Ungaro R, Abouhamze A, Moldawer LL. 2007. INCREASED MORTALITY AND ALTERED IMMUNITY IN NEONATAL SEPSIS PRODUCED BY GENERALIZED PERITONITIS. Shock 28:675–683. doi:10.1097/shk.0b013e3180556d09

Zantl N, Uebe A, Neumann B, Wagner H, Siewert J-R, Holzmann B, Heidecke C-D, Pfeffer K. 1998. Essential Role of Gamma Interferon in Survival of Colon Ascendens Stent Peritonitis, a Novel Murine Model of Abdominal Sepsis. Infect Immun 66:2300–2309. doi:10.1128/IAI.66.5.2300-2309.1998

